# Genomic Heritability: A Ragged Diagonal Between Bias and Variance

**DOI:** 10.1101/2021.09.19.460999

**Authors:** Mitchell J. Feldmann, Hans-Peter Piepho, Steven J. Knapp

**Author notes:** Corresponding author: Department of Plant Sciences, University of California, Davis, California, United States of America.

## Abstract

Many important traits in plants, animals, and microbes are polygenic and are therefore difficult to improve through traditional marker-assisted selection. Genomic prediction addresses this by enabling the inclusion of all genetic data in a mixed model framework. The main method for predicting breeding values is genomic best linear unbiased prediction (GBLUP), which uses the realized genomic relationship or kinship matrix (**K**) to connect genotype to phenotype. The use of relationship matrices allows information to be shared for estimating the genetic values for observed entries and predicting genetic values for unobserved entries. One of the key parameters of such models is genomic heritability 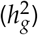, or the variance of a trait associated with a genome-wide sample of DNA polymorphisms. Here we discuss the relationship between several common methods for calculating the genomic relationship matrix and propose a new matrix based on the average semivariance that yields accurate estimates of genomic variance in the observed population regardless of the focal population quality as well as accurate breeding value predictions in unobserved samples. Notably, our proposed method is highly similar to the approach presented by Legarra (2016) despite different mathematical derivations and statistical perspectives and only deviates from the classic approach presented in VanRaden (2008) by a scaling factor. With current approaches, we found that the genomic heritability tends to be either over- or underestimated depending on the scaling and centering applied to the marker matrix (**Z**), the value of the average diagonal element of **K**, and the assortment of alleles and heterozygosity (*H*) in the observed population and that, unlike its predecessors, our newly proposed kinship matrix **K**_*ASV*_ yields accurate estimates of 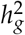 in the observed population, generalizes to larger populations, and produces BLUPs equivalent to common methods in plants and animals.

## Introduction

Linear mixed model (LMM) analyses are routine in breeding and quantitative genetics across organisms and are used in the prediction of breeding values in plants and animals (Henderson 1977; VanRaden 2008; Hayes *et al*. 2009; Albrecht *et al*. 2011; Endelman 2011; Crossa *et al*. 2014; Meuwissen *et al*. 2016)and polygenic risk scores (PRSs) in humans (de los Campos *et al*. 2010; Makowsky *et al*. 2011; Dudbridge 2013; Maier *et al*. 2018; Wray *et al*. 2019; Truong *et al*. 2020), partitioning of sources of variance (Searle *et al*. 1992; Lynch and Walsh 1998; Visscher *et al*. 2008; Kang *et al*. 2010; Piepho *et al*. 2012a; Schmidt *et al*. 2019; Feldmann *et al*. 2021), and controlling for confounding effects in genome-wide association studies (GWAS) (Yu *et al*. 2006; Visscher *et al*. 2012; Korte and Farlow 2013; Visscher *et al*. 2017). Genomic prediction approaches are widely applied in the study of complex traits in natural and experimental populations and facilitate the estimation of genomic variance 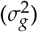, genomic heritability 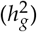, and other quantitative genetic parameters (Meuwissen *et al*. 2001; Crossa *et al*. 2010; de los Campos *et al*. 2015; Lehermeier *et al*. 2017; Wray *et al*. 2019) and has been widely adopted in plant breeding, human genetics, and biology (Meuwissen *et al*. 2001; Habier *et al*. 2007; Goddard and Hayes 2007; Heffner *et al*. 2009; Bloom *et al*. 2013). These methods yield estimates of genomic-estimated breeding values (GEBVs) for both phenotyped and unphenotyped individuals, in addition to estimates of genomic parameters, e.g., genomic variance (VanRaden 2008; Goddard 2009; Forni *et al*. 2011; de los Campos *et al*. 2015).

When applying genomic prediction, the genetic variance in the training population can be estimated using ‘ genomic variance’ 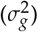—the genetic variance explained by genome-wide associations between the underlying QTL and DNA markers genotyped in the training population (Meuwissen *et al*. 2001; Habier *et al*. 2013; de los Campos *et al*. 2015). Genomic variance is often estimated from LMM analyses using a genomic relationship matrix (GRM), e.g., GBLUP, which is calculated from SNP markers to measure the relatedness among entries (VanRaden 2008; Yang et al. 2010; Habier et al. 2013). Genomic variance is found in many ratios throughout modern quantitative genetic research, including: genomic heritability, predictive ability, selection reliability, prediction error variance, and response to genomic selection (Goddard 2009; Hickey *et al*. 2009; Crossa *et al*. 2014; Gorjanc *et al*. 2015; de los Campos *et al*. 2015). Of these ratios, genomic heritability has been the most frequently reported in studies.

Genomic heritability is defined as:

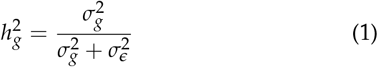

where 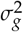 is the genomic variance and 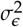 is the residual variance on an entry-mean basis and is typically estimated by substituting REML variance component estimates (VCEs) into (1). We studied how different forms of **K** affect VCEs and found that there are systematic differences in the VCEs arising from different forms of **K** (VanRaden 2008; Astle *et al*. 2009; Yang *et al*. 2010; Forni *et al*. 2011; Endelman and Jannink 2012; Legarra 2016) and the resulting VCEs may not always be accurate when directly substituted into (1), as is common practice, to obtain genomic heritability estimates. Specifically, genomic heritability will be either over- or underestimated when REML variance estimates associated with different forms of **K** are substituted into (1) when the observed population is not at HWE; however, as Strandén and Christensen (2011) clearly articulated, the GEBVs are equivalent in studies that employ different methods of calculating **K** and different allele coding for **Z** (Powell *et al*. 2010). Despite this, researchers are using the same approaches for both prediction and estimation simultaneously, and may be reporting incorrect genomic heritability estimates.

Here, using the average semivariance approach (Piepho 2019; Feldmann *et al*. 2021), we derive a new form for **K**, referred to as **K**_*ASV*_, which is calculated as the product 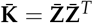 from the mean-centered marker matrix 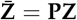, where 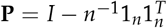 is the idempotent, mean-centering *n* × *n*-matrix (VanRaden 2008; Endelman and Jannink 2012). The average semivariance relationship matrix is:

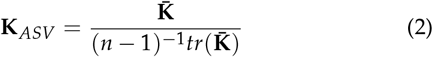

This matrix is scaled to the residual variance-covariance matrix and directly yields accurate estimates of 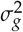 and 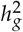. It is possible to scale other forms of the GRM by dividing **K** by (*n* − 1)^−1^*tr*(**K**) and to scale estimates of genomic variance from any form of **K** by multiplying (*n* − 1)^−1^*tr*(**K**) by 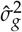 to obtain average semivariance VCEs. We explore the practical implications of **K**_*ASV*_ for estimating 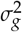 and 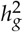 in a wild population of *Arabidopsis thaliana* (Atwell *et al*. 2010), a wheat (*Triticum aestivium*) breeding population (Crossa *et al*. 2010), a laboratory mouse (*Mus musculus*)population (Valdar *et al*. 2006), an apple (*Malus* × *domestica*) breeding population (Kumar *et al*. 2015), and a pig (*Sus scrofa*) breeding population (Cleveland *et al*. 2012).

## Results and Discussion

### Average semivariance directly yields accurate estimates of genomic variance and genomic heritability

The average semivariance (ASV) estimator of total variance (Piepho 2019; Schmidt *et al*. 2019; Feldmann *et al*. 2021) is half the average total pairwise variance of a difference between entries and can be decomposed into independent sources of variance, e.g., genomic and residual. There are two alternative ASV derivations, both leading to the same definitions of the estimators. The first derivation originated in geostatistics and estimates the semivariance among entries *i* and *j* or half of the variance among all pairwise differences among observations: 2^−1^*var*(*y_i_* − *y_j_*) (Webster and Oliver 2007; Piepho 2019). The second derivation utilizes the sample variance: 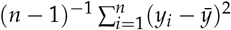 and its expectation, as in Estaghvirou *et al*. (2013). The analyses shown throughout this paper assume the dependent variables are best linear unbiased estimators (BLUEs) or least square means (LSMs) for entries (*y*). The residual variance of the LSMs is given by 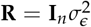. The average semivariance can easily deal with more general forms of variance-covariance matrices in generalized LMMs (Piepho 2019).

The LMM for this analysis is:

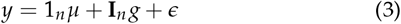

where *y* is the vector of phenotypic LSMs of for *n* entries, *n* is the number of entries, 1_*n*_ is an *n*-element vector of ones, *μ* is the population mean, **I**_*n*_ is the identity matrix of size *n*, *g* is an *n*-element vector of random effect values for entries with 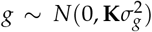, and *ϵ* is the residual for the each entry with 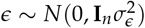.

In the `sommer` R package LMM (3) is expressed as:

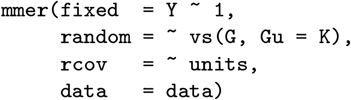

where `data` is an *n* × 2 matrix with `Y` as a column of LSMs and `G` is a column of factor coded entry IDs (Covarrubias-Pazaran 2016).

The ASV definition of total variance from LMM (3) is:

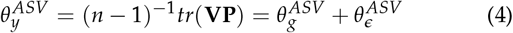

where 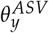 is the total phenotypic variance, 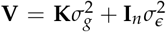 is the variance covariance among observations, 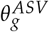 is the genomic average semivariance, and 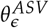 is the average semivariance of the residuals.

The ASV definition of the genomic variance is:

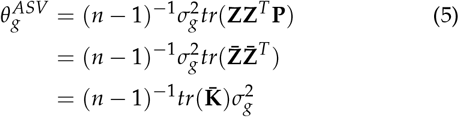

where 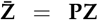 is the mean-centered marker matrix, and 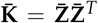 is the realized genomic relationship or kinship matrix described by VanRaden (2008) omitting the scaling constant 2∑_*j*_ *p_j_* (1 − *P_j_*), where *p_j_* is the allele frequency of the *j*-th SNP, which requires HWE to hold (de los Campos *et al*. 2015), and 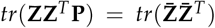. The trace of 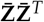 is a function of heterozygosity in the observed population (Vitezica *et al*. 2013, 2017; Legarra *et al*. 2018). When the observed population is in HWE, 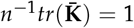, and when the population is not in HWE due to inbreeding, the 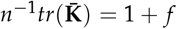, where *f* is the in coefficient of inbreeding (Endelman and Jannink 2012; Legarra *et al*. 2018).

In the general case, 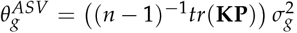, where **K** is any form of the genomic relationship matrix calculated from **Z**, without centering, or 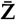, with centering, because 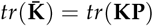. If we assume 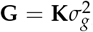, where **G** is the variance-covariance of GEBVs, it can observed the magnitude *tr*(**K**) is directly inverse to the magnitude of 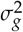, assuming that 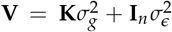 and that 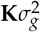 and 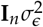 are uncorrelated. Thus, as different scales are applied to **K**, the inverse should be applied to 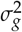.

The ASV definition of the residual variance is:

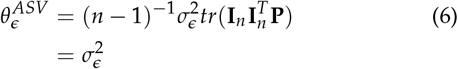

Importantly, the genomic variance 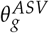 is on the same scale as the residual variance 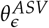 and both are defined such that (4) is true. Note that REML estimates are directly equivalent to ASV estimates when BLUEs or LSMs are used as the response variable *y*.

Substituting (5) and (6) into (1), the ASV estimator of genomic heritability 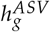 is:

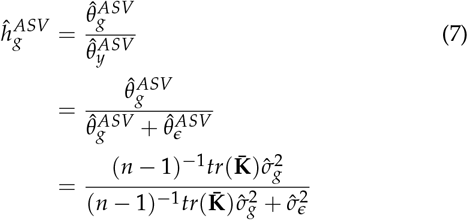

This formulation can be used directly with any form of **K** or 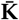 by substitution of REML VCEs.

There are two equivalent solutions for obtaining ASV estimates of 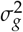. We can either correct the genomic and error variances from (7) using any form of **K** or directly acquire these corrected estimates by using an alternative form of the genomic relationship matrix given in (2). Note that the denominator 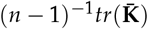 is a scalar and is derived from the sample size and the covariance among the observed individuals. **K**_*ASV*_ estimates genomic variance on the same scale as the residual error variance (Xu 2013; Vitezica *et al*. 2017), which is essential for accurately calculating genomic heritability and comparing across populations (Legarra 2016). We reiterate and reemphasize that the mean of the diagonal elements of **K** is 1 and the mean element of **K** is 0 in order to compare genomic heritability in human, plant, and animal genetics across populations and studies (Hayes *et al*. 2009; Speed and Balding 2015; Legarra 2016).

### Analysis of simulated data confirms that ASV yields accurate VCEs

The ASV relationship matrix yielded accurate estimates of genomic heritability in the observed populations, while the other methods varied with the level of heterozygosity. When heterozygosity *H* < 0.5 the genomic variance tends to be underestimated and when *H* > 0.5 the genomic variance tends to be overestimated (Fig 1). This was true regardless of the population size; e.g., *n* = 250, 500, and 1, 000. All methods tend to produce accurate estimates when *H* = 0.5, in which case the inbreeding coefficient *f* = 0 and HWE is not violated.

**Figure 1.**
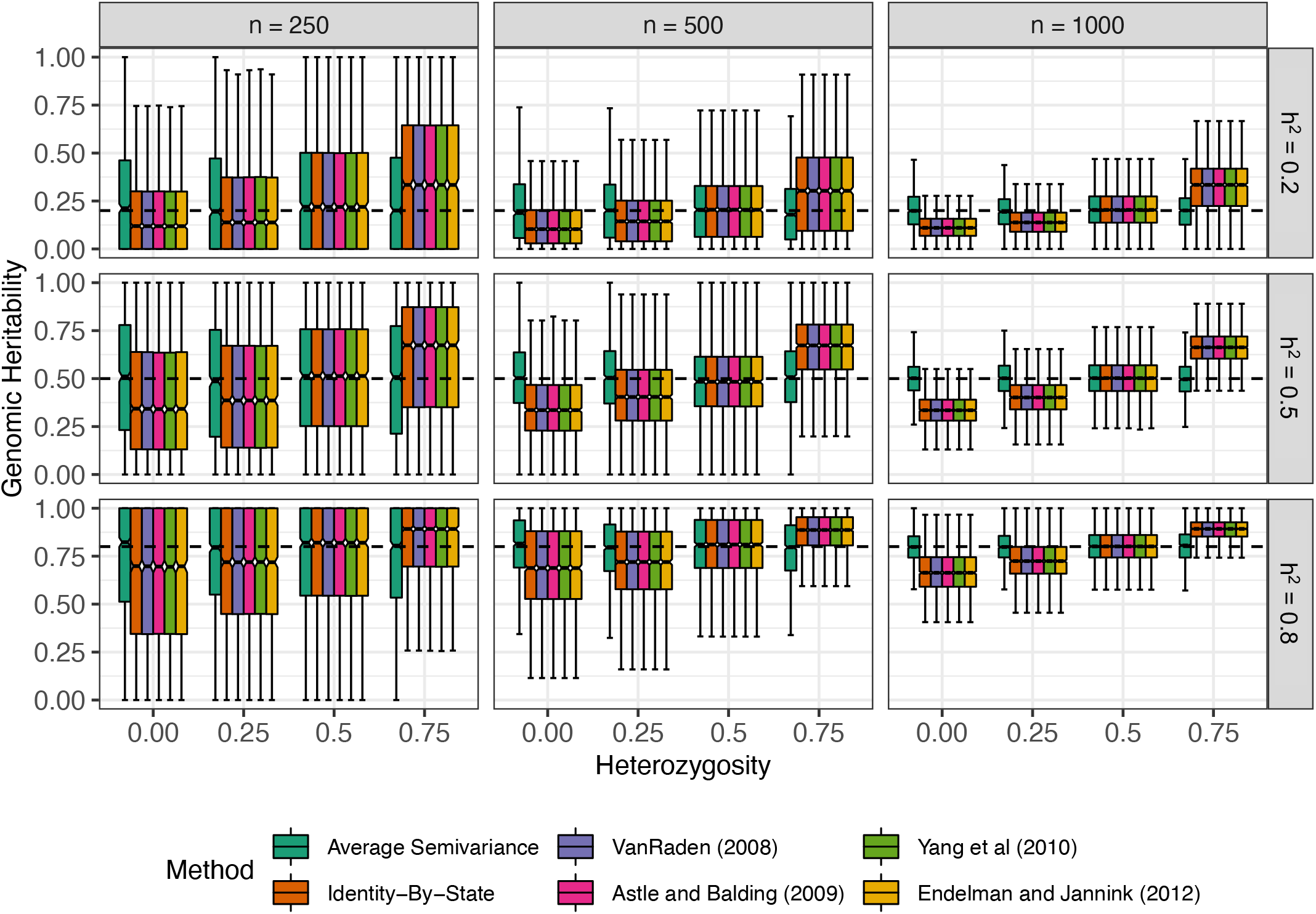
Effect of heritability 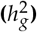, population size (*n*), and heterozygosity (*H*) on the accuracy of genomic heritability estimates. Phenotypic observations were simulated for 1,000 samples with *n* = 250, 500, and 1000 (left to right) genotyped for *m* = 5, 000 SNPs and the average heterozygosity *H* = 0, 25, 50, and 75%. The accuracy of genomic heritability estimates 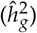 from the six relationship matrices is shown for true genomic heritability 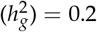 (upper panel), 0.5 (middle panel), and 0.8 (lower panel). The upper and lower halves of each box correspond to the first and third quartiles (the 25th and 75th percentiles). The notch corresponds to the median (the 50th percentile). The upper whisker extends from the box to the highest value that is within 1.5 · *IQR* of the third quartile, where *IQR* is the inter-quartile range, or distance between the first and third quartiles. The lower whisker extends from the first quartile to the lowest value within 1.5 · *IQR* of the quartile. The dashed line in each plot is the true value from simulations.

The precision (variance) improved by increasing the population size (*n*), but the accuracy (bias) did not improve. It has been demonstrated *ad nauseam* that increasing *n* increases precision, or lowers the sample variance of the estimates, but does not eliminate bias (Laird and Ware 1982; Searle *et al*. 1992; Lynch and Walsh 1998). Notably, the entire parameter space of 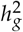 is observed when the population size is small (Fig 1). The standard deviation of the estimates of 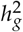 depends the sample size *n* and the true value of 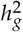, (Fig 1). Only the ASV approach yields stable precision as *H* increases and all other methods that we examined have variable precision, as well as variable accuracy at different values of *H* (Fig 1).

Inbreeding changes the pattern of genomic variance to be completely among individuals and drives deviations from HWE (Bernardo 2002). In inbred plant species, e.g., maize, rice, wheat, sorghum, it is common practice to estimate the genomic variance from LMM (3) and then to scale the estimated variance component by a factor of 2 in both the numerator and denominator of (1) (Cockerham 1983; Wricke and Weber 2010; Bernardo 2002; Isik *et al*. 2017). In RILs and doubled haploids (DHs), the standard GRMs, excluding **K**_*ASV*_ and **K**_*GN*_, have *n*^−1^*tr*(**K**) = (1 + *f*) = 2 (Endelman and Jannink 2012). This factor of 2 is, in fact, the same factor of 2 classically used to scale the VCEs in inbred plant species (Cockerham 1983; Wricke and Weber 2010; Bernardo 2002; Isik *et al*. 2017). Extending this implies that there is a scaling factor for intermediate levels of inbreeding, e.g., − 1 ≤ *f* ≤ 1, not just when *f* = 1, as in RILs and DHs, which is directly accounted for **K**_*ASV*_.

Experiment designs that enable screening of a greater number of entries *n* may help researchers obtain more precise estimates of key parameters in resource restricted programs (Smith *et al*. 2006; Moehring *et al*. 2014; Mackay *et al*. 2019; Borges *et al*. 2019; Hoefler *et al*. 2020) and ASV can ensure that those estimates are accurate and comparable across populations (Legarra 2016). In many plant quantitative genetic studies, the population sizes are *n* ≈ 500, which may pose a general problem for variance component and ratio estimation as those variance components tended to have high variability between replicated experiments (Fig 1). For very large populations, common in human and domesticated animal studies, it is possible to imagine precise (low variance) but inaccurate (high bias) estimates of 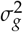 and 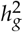 resulting from different relationship matrices, unless the assumptions of HWE happen to be perfectly met in the study population.

### Analysis of real-world data confirms that ASV does not impact BLUPs

The differences in the prediction accuracy for unobserved individuals in the examples we selected, when present, appeared to be negligible and did not lend themselves clearly to “better” or “worse” categorisations (Fig 2). Strandén and Christensen (2011) indicated, and our analyses confirmed, that different centering and scaling methods of markers do not greatly affect the accuracy of genomic prediction as demonstrated in Fig 2 for five different species and 22 different traits. The elements of **K**_*ASV*_ are strongly correlated to the elements of **K**_*VR*_. **K**_*VR*_ is widely used in genomic prediction and selection literature in plants and animals (Fig 2), which suggest that **K**_*ASV*_ could be easily integrated into existing pipelines and analyses. This is to be expected, as the accuracy of genomic predictions is invariant to marker coding, centering, or scaling of **K** (Powell *et al*. 2010; Strandén and Christensen 2011). Any form of **K** can be used to estimate *g* and *β*—the random marker effects— because ridge regression is a linear transformation of LMM (3), e.g., 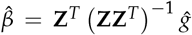 (Piepho 2009; Piepho *et al*. 2012b; Morota and Gianola 2014). Thus, for genomic selection applications, researchers can apply whichever form of the realized relationship matrix they choose to obtain GEBVs 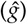 and estimated random marker effects 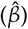 and to calculate prediction accuracy 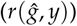.

**Figure 2.**
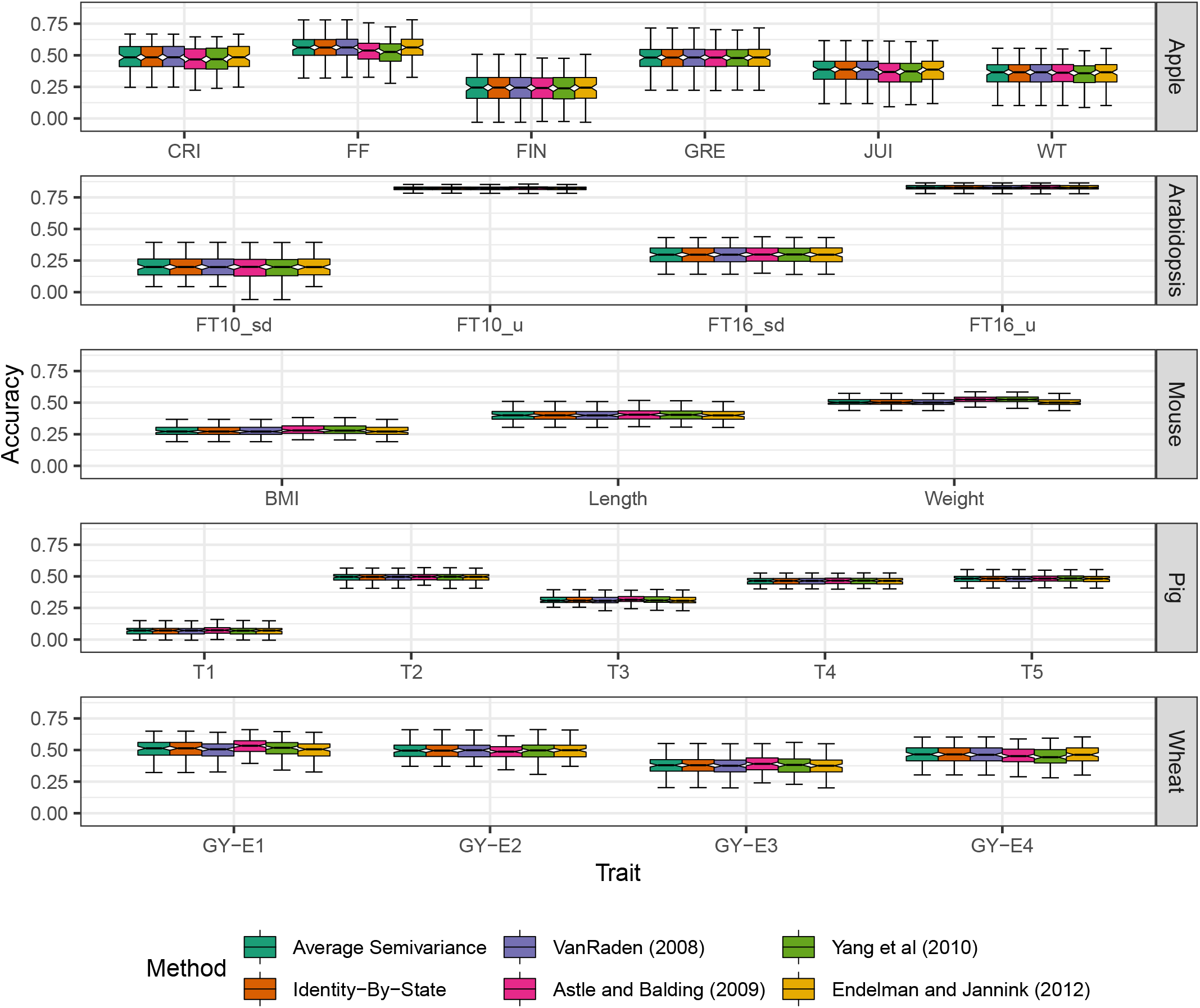
Cross-validated prediction accuracy for different species and traits using different genomic relationships. Results are presented from 100 realizations of 80 : 20 cross-validation using the six relationship matrices for six traits in an apple population with *n* = 247 entries genotyped at *m* = 2, 829 SNPs (Kumar *et al*. 2015) (first row), four traits in an Arabidopsis population with *n* = 1, 057 entries genotyped at *m* = 193, 697 SNPs (Atwell *et al*. 2010) (second row), three traits in an mouse population with *n* = 1, 814 entries genotyped at *m* = 10, 346 SNPs (Valdar *et al*. 2006) (third row), and five traits in a pig population with *n* = 3, 534 entries genotyped at 52, 843 SNPs (Cleveland *et al*. 2012) (fourth row), four traits in an wheat population with *n* = 599 entries genotyped at *m* = 1, 278 SNPs (Crossa *et al*. 2010) (fifth row). For the Arabidopsis data set (second row), **K**_*ϒ*_ systematically produced singular systems in `sommer::mmer()` and prediction accuracy was not estimated for either *FT*10_*μ*_ or *FT*16_*μ*_. The upper and lower halves of each box correspond to the first and third quartiles (the 25th and 75th percentiles). The notch corresponds to the median (the 50th percentile). The upper whisker extends from the box to the highest value that is within 1.5 · *IQR* of the third quartile, where *IQR* is the inter-quartile range, or distance between the first and third quartiles. The lower whisker extends from the first quartile to the lowest value within 1.5 · *IQR* of the quartile.

While the choice of **K** does not impact BLUP, the choice of GRM impacts estimates of genomic variance 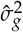, genomic heritability 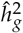, predictive ability 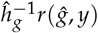, average prediction error variance *PEV*, and selection reliability 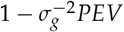, which all rely on 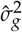. The *PEV* of the BLUPs was similar across relationships indicating that differences in selection reliability and predictive ability will be driven by differences in 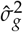 that arise from different forms of **K**. These results are also preceded by Strandén and Christensen (2011) who show that *PEV* depends on the centering and scaling and that different methods change the amount of certainty, or reliability, in the BLUPs.

We observed the expected patterns that were exposed by our simulations (Fig 1) in the real-world data sets (Table 1). When the study populations are fully inbred as in wheat, Arabidopsis, or inbred *per se* evaluations in hybrid crops, such as maize, tomato, rice, the ASV corrections produce larger VCEs (Table 1). Similarly, for populations where *H* ≈ 0.5 the relationship matrices all tend to have the average diagonal element of **K** equal to 1 and thus the scale of genomic and residual variances are similar (Xu 2013; Legarra 2016; Vitezica *et al*. 2017). Our analyses of the flowering time data yielded equivalent results and patterns to Lehermeier *et al*. (2017) suggesting that **K**_*ASV*_ may be providing an estimate of genomic variance that naturally accounts for LD (Table 1). However, the empirical similarities in the Arabidopsis case study between our approach and Lehermeier *et al*. (2017)’s method M2 strongly implicates a non-circumstantial link and that ASV estimates of genomic variance may naturally account for LD.

**Table 1.**
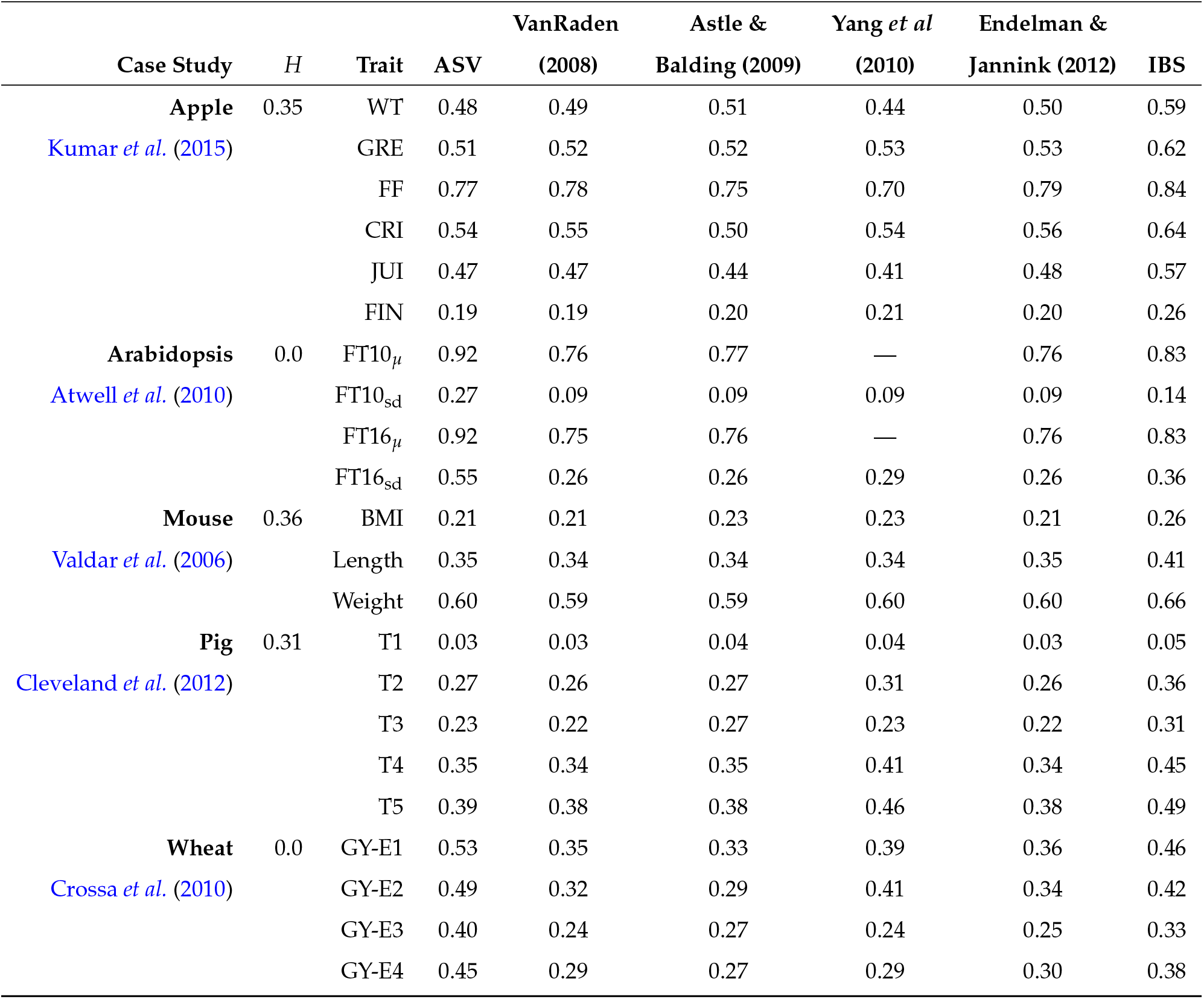
Genomic heritability 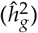 estimates for six traits in an apple population with *n* = 247 entries genotyped at *m* = 2, 829 SNPs (Kumar *et al*. 2015), four traits in an wheat population with *n* = 599 entries genotyped at *m* = 1, 278 SNPs (Crossa *et al*. 2010), four traits in an Arabidopsis population with *n* = 1, 057 entries genotyped at *m* = 193, 697 SNPs (Atwell *et al*. 2010), and three traits in an mouse population with *n* = 1, 814 entries genotyped at *m* = 10, 346 SNPs (Valdar *et al*. 2006), and five traits in a pig population with *n* = 3, 534 entries genotyped at 52, 843 SNPs (Cleveland *et al*. 2012) using the six methods compared in this article.

### The relationship between ASV and normalized genomic relationship matrices

As our simulations confirmed, the scaling of the trace of **K** profoundly affects the estimation of 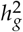 (Figure 1), consistent with Speed and Balding (2015) and Legarra (2016), but has virtually no effect on BLUP or genomic prediction accuracy (Fig 2). Speed and Balding (2015) argued that the GRM, **K**, should be normalized such that 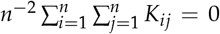 and *tr*(**K**) = *n*, where *n* is the number of entries, e.g., individuals, families, or strains, and *K_ij_* element in the *i*-th row and *j*-th column of **K** (Forni *et al*. 2011; Legarra *et al*. 2015; Legarra 2016). The rationale for their proposal was more fully articulated by Legarra (2016); however, the mathematical conditions Speed and Balding (2015) proposed are seldom met in practice because of inbreeding, LD, and deviations from Hardy-Weinberg equilibrium (HWE). Further, we showed that ASV yields unbiased estimates of 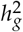 that are invariant to the heterozygosity (*H*) of markers in the population, while other methods are strongly affected by *H* and yield biased VCEs when *H* ≠ 0.5 (Fig 1)(Legarra *et al*. 2018). We found that the normalized **K**, i.e., **K**_*GN*_, proposed by Forni *et al*. (2011) (discussed by Legarra (2016)) yields estimates of **K** that are virtually identical to ASV (**K**_*ASV*_). Although these estimators were found through different approaches, we show here that they are mathematically equivalent apart from a single degree of freedom difference in the divisor of the GRM: Legarra (2016) used the number of observations (*n*), whereas we used *df_g_* = *n* − 1 for calculating the sample variance (Bulmer 1979). Here, we demonstrate the similarities between **K**_*ASV*_ and **K**_*GN*_.

Recall that the unbiased estimator of a sample variance among breeding values is:

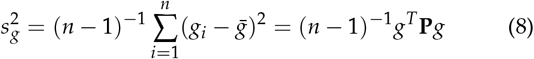

where 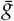 is the average genetic value of individuals in the observed population and *g_i_* is the genetic value of the *i*-th observed individual. The population variance, which is considered by Legarra (2016), among breeding values is:

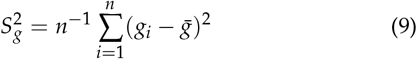

We use 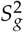, the population variance, to indicate a different metric relative to 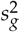, the sample variance.

Taking expectations of 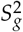 with respect to the distribution of *g*:

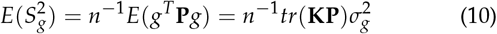

Thus, the genomic variance in the reference population 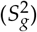 will be a function of the variance component associated with **K** in the mixed model (Legarra 2016). While Legarra (2016) does not make an explicit statement about how **K** could be formulated, it is clear that a genomic relationship could be defined as:

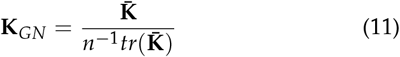

This form was explicitly introduced as the normalized relationship matrix by Forni *et al*. (2011) and has been used in several studies. There are likely to be differences between these **K**_*ASV*_ and **K**_*GN*_ when *n* is small. When *n* is reasonably large **K**_*ASV*_ ≈ **K**_*GN*_ as *n* → ∞ (Fig 1). This work, Forni *et al*. (2011), and Legarra (2016) both arrive at numerically similar solutions through conceptually different derivations, which we feel is indicative of the value of these approaches for plant, animal, and human genetic studies that rely on genomic relatedness, e.g, genome-wide association studies, genomic prediction, or inferring population structure and ancestry.

## Conclusions

The interpretation of genomic variance and heritability estimates was shown to be affected by the methods used to estimate **K**. We provide a relationship matrix **K**_*ASV*_ that directly yields consistent VCEs and a correction factor (*n* − 1)^−1^*tr*(**K**) that will allow users to obtain accurate estimates of genomic heritability in the observed population using various software packages (Misztal *et al*. 2002; Clifford and McCullagh 2006; Vazquez *et al*. 2010; Endelman 2011; Zhou and Stephens 2012; Pérez and de Los Campos 2014; Akdemir and Okeke 2015; Covarrubias-Pazaran 2016; Bürkner 2017; Runcie and Crawford 2019; Caamal-Pat *et al*. 2021; Butler 2021). ASV is a strategy that can be used for estimating and partitioning the total variance into component parts (Piepho 2019), such as the variance explained by large effect markers and marker-marker interactions (Feldmann *et al*. 2021). As advocated by Speed and Balding (2015) and Legarra (2016), the ragged diagonal elements of **K**_*ASV*_ equal 1, on average, and the off-diagonal elements equal, on average. ASV directly yields accurate estimates of genomic heritability in the observed population and can be used to adjust deviations that arise from other commonly used methods for calculating genomic relationships regardless of the population constitution, such as inbred lines and F_1_ hybrids, unstructured GWAS populations, and animal herds or flocks (Fig 1). The ASV approach to calculate the realized relationship matrix has the property of unit diagonals and a mean value of zero (Speed and Balding 2015; Legarra 2016), which are both necessary for accurate VCEs. We believe that **K**_*ASV*_ provides a powerful approach for directly estimating genomic heritability for the observed population regardless of study organism or design. In conclusion, our recommendation is that the average semivariance approach be considered for adoption by genetic researchers working in humans, microbes, or (un)domesticated plants and animals.

## Methods and Materials

### Computer Simulations

We generated 36 experiment designs with different heterozygosity *H* = 0.0, 0.25, 0.5, and 0.75 and different trait heritability 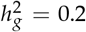, 0.5, and 0.8 and for population sizes of *n* = 250, 500, and 1, 000. In all examples, 1, 000 populations genotyped at *m* = 5, 000 causal loci were used to generate the genetic traits. We simulated marker effects for all *m* = 5, 000 loci following a normal distribution *μ* = 0 and *σ* = 1. When multiplied by the marker genotypes and summed, the score is taken as the true genetic value *g* of each individual. Residuals are simulated with *μ* = 0 and 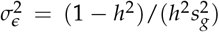 to obtain a trait with the desired genomic heritability (Endelman 2011) and 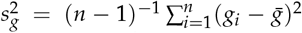 is the sample variance among genotypic values (Estaghvirou *et al*. 2013). All plots are made with the `ggplot2` package (Wickham 2016) in R 4.0.2 (R Core Team 2020).

For each population, simulated or case example, we calculated and applied six relationship matrices, including **K**_*ASV*_. We used `AGHmatrix::Gmatrix()` to calculate the Yang *et al*. (2010) (**K**_*ϒ*_) and VanRaden (2008) relationship (**K**_*VR*_) matrices (Rampazo Amadeu *et al*. 2016), `rrBLUP::A.mat()` to calculate the Endelman and Jannink (2012) (**K**_*EJ*_) relationship matrix, and `statgenGWAS::kinship()` to estimate the Astle *et al*. (2009) (**K**_*AB*_) and IBS relationship (**K**_*IBS*_) matrices (van Rossum and Kruijer 2020).

The form proposed by VanRaden (2008) is:

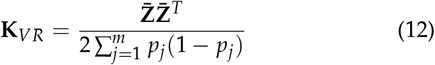

where 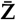 is the marker matrix centered on column means (2*p_j_*), and *p_j_* is the minor allele frequency for the *j*-th SNP. This form assumes HWE and that *p_j_* is obtained from a historical reference population, and not the observed population. When *p_j_* is estimated in the population of observed entries, the centering by 2*p_j_* is equivalent to column centering and **K**_*VR*_ only differs from **K**_*ASV*_ by a scaling factor, such that: **K**_*ASV*_ × ((*n* − 1)^−1^*tr*(**K**_*VR*_)) = **K**_*VR*_ and **K**_*VR*_/ ((*n* − 1)^−1^*tr*(**K**_*VR*_)) = **K**_*ASV*_.

The form of the relationship matrix proposed by Endelman and Jannink (2012) is :

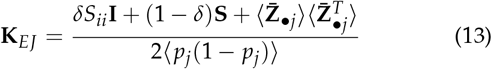

where *δ* ≈ (*n/m*)*CV*^−2^ is a shrinkage factor, *CV*^2^ is the coefficient of variation of the eigenvalues of **S**, 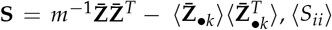, 〈*S_ii_*〉 is the mean of diagonal elements of **S**. Notably, at high marker densities, when *δ* = 0,Endelman and Jannink (2012) is equivalent to VanRaden (2008) when *p_j_* is estimated from the observed entries.

The method proposed by Yang *et al*. (2010) also centers the columns of **Z** by subtracting 2*p_j_*

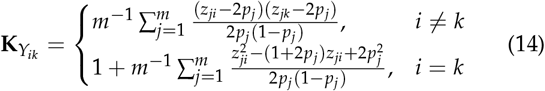

where *Z_ij_* is the *j*-th SNP in the *i*-th individuals, *Z_jk_* is the *j*-th SNP in the *k*-th individual when *j* ≠ *k*, and *m* is the number of markers. It is clear that the diagonals are treated differently, relative to the off-diagonals.

The method proposed by Astle *et al*. (2009) is:

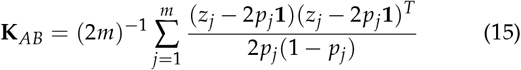

where *Z_j_* is the *i*-element vector of the *j*-th SNP.

The classical identity-by-state definition is given in Astle *et al*. (2009) as:

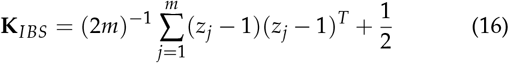

Note that this is the the only calculation that is not scaled or centered by any function of *p_j_*. In general, the IBS form is most similar to Astle *et al*. (2009), while VanRaden (2008) and Endelman and Jannink (2012) are equivalent when the marker density is high and the study population is considered as the base population.

For each model and each simulation, we estimated two variance components (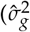 and 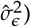) using `sommer::mmer()` and took the ratio of variance components from these models using `sommer::vpredict()` in R (R Core Team 2020). We estimated genomic heritability using the standard form by substituting REML estimates from (3) into (1). In this study, 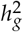 was either 0.2, 0.5, or 0.8.

### Example Data

We analyzed four publicly available data sets using six methods for calculating the realized relationship matrix and estimated 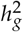. First, we analyzed six traits from Kumar *et al*. (2015) which evaluated a breeding population of *n* = 247 apple (*Malus* × *domestica*) hybrids genotyped a *m* = 2, 829 SNPs with *H* = 0.348 (Kumar *et al*. 2015). The reported traits were fruit weight (WT), fruit firmness (FF), greasiness (GRE), crispiness (CRI), juiciness (JUI), and flavor intensity (FIN). The shrinkage factor *δ* from Endelman and Jannink (2012) was equal to 0.02. Second, we analyzed the wheat data set from Crossa *et al*. (2010) which evaluated *n* = 599 wheat (*Triticum aestivum*) fully inbred lines (*H* =0.0; *δ* = 0.03) for grain yield (GY) in four environments genotyped for *m* = 1, 278 SNPs. In this paper, we evaluated each environment, GY-E1, GY-E2, GY-E3, and GY-E4, with an independent model. Third, we analyzed data from Valdar *et al*. (2006) which evaluated a laboratory population of *n* = 1, 814 stock mice (*Mus musculus*) for body mass index (BMI), body length, and weight and genotyped for *m* = 10, 346 SNPs (*H* =0.363; *δ* = 0.01). Fourth, we analyzed a population of *n* = 1, 057 naturally occurring Arabidopsis (*Arabidopsis thaliana*) ecotypes from Atwell *et al*. (2010) and Alonso-Blanco *et al*. (2016) phenotyped for mean (*μ*) and standard deviation (sd) of flowering time under 10° C (FT10) and 16° C (FT16) and genotyped at *m* = 193, 697 SNPs (*H* =0.0; *δ* = 0.0). Fifth, we analyzed a commercial pig (*Sus scrofa*) population made available by PIC (a Genus company) with *n* = 3, 534 entries genotyped at *m* = 52, 843 SNPs (*H* =0.311; *δ* = 0.0) that were phenotyped for five traits: T1, T2, T3, T4, and T5 (Cleveland *et al*. 2012). For each population, we calculate the 6 relationship matrices (12–16) and apply them in (3) for each trait to estimate 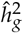 with (1). Similarly, we used `sommer::mmer()` to estimate the prediction error variance (*PEV*) of BLUPs by setting the option *getPEV=T* (Covarrubias-Pazaran 2016; Yu and Morota 2021).

We performed cross-validation to determine prediction accuracy 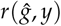. We first split each population in 80% train and 20% test and estimated genomic BLUPs and then calculated the accuracy as the correlation between the estimated LSM *y* and the BLUP 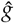 for all entries in the test set. We performed this cross-validation scheme 100 times for each population and each trait.

## Data Availability

The input and output from simulations and analyses has been deposited in a public Zenodo repository (https://doi.org/10.5281/zenodo.5038485).

## Acknowledgments

The authors thank Dr. Andres Legarra for review suggestions that greatly improved the manuscript.

## Author Contributions

**Conceptualization:** MJF, HPP, SJK **Data curation:** MJF **Formal Analysis:** MJF **Funding Acquisition:** HPP, SJK **Investigation:** MJF, HPP, SJK **Methodology:** MJF **Project administration:** MJF, HPP, SJK **Resources:** MJF, HPP, SJK **Software:** MJF **Supervision:** MJF, HPP, SJK **Validation:** MJF **Visualization:** MJF **Writing – original draft preparation:** MJF, HPP, SJK **Writing – review & editing:** MJF, HPP, SJK

